# Significant Broad Spectrum Antiviral activity of Bi121 Against Different Variants of SARS-CoV-2

**DOI:** 10.1101/2022.01.29.477140

**Authors:** Bobban Subhadra, Ragini Agrawal, Virender Kumar Pal, Agnes-Laurence Chenine, Jeffy George Mattathil, Amit Singh

## Abstract

The SARS-CoV-2 pandemic infected 343 million people with over 5.59 million deaths. New mutated lineages of SARS-CoV-2 such as Omicron are evolving faster. Broad-spectrum viral inhibitors that block the initial stage of infection by reducing virus proliferation and disease severity is an unmet global medical need. We studied Bi121, a standardized polyphenolic-rich compound isolated from *Pelargonium sidoides*, against recombinant Vesicular Stomatitis Virus (rVSV)-pseudotyped SARS-CoV-2S (spike) that represent mutations in the spike protein of six different variants of SARS-CoV-2. Bi121 was effective in neutralizing all six rVSV-ΔG-SARS-CoV-2S variants expressing different mutations. The antiviral activity of Bi121 was then assessed against three variants of SARS-CoV-2 (USA WA1/2020, Hongkong/VM20001061/2020, B.1.167.2 (Delta)) using RT-qPCR and plaque assays in two different cell lines (Vero cells and HEK-ACE2). Bi121 showed significant activity toward all the three variants tested, suggesting a broad-spectrum activity.

## Introduction

The SARS-CoV-2 pandemic infected 343 million people with over 5.59 million deaths, markedly affecting human life and existence (1). In the US alone, over 883,903 deaths have been reported as of January 24, 2022 (2). Since its first report, several SARS-CoV-2 lineages evolved in the last 24 to 30 months (3); currently, the Omicron variant is dominant around the world, but researchers are monitoring other variants of concern, including the recently discovered variant of Omicron which was reported in Denmark. Evolutionary viral mutations in spike and other proteins in highly evolved lineages may warrant survival benefits to the virus to thwart the human immune system (4). Omicron variant carries 46 high prevalence mutations specific to Omicron; Twenty-three of these are localized within the spike (S) protein and the rest localized to the other three viral structural proteins of the virus (5).

PAXLOVID, in combination with ritonavir, developed by Pfizer, was found to reduce the risk of hospitalization or death by 89% compared to placebo in non-hospitalized high-risk adults with SARS-CoV-2 (6). Because of the prolific nature of viral infectivity and evolutionary pressure from vaccine-induced adaptive immune response and prospective antiviral treatments, there is a high probability of the emergence of more infectious SARS-CoV-2 variants with potential drug resistance (7). This suggests the need for antivirals with broad-spectrum neutralization activity towards various lineages of existing and future SARS-CoV-2 variants.

Several single compound antivirals have shown to be effective against SARS-CoV-2 in preclinical studies and are in clinical development. However, these single molecule-single target-based approaches may be challenging in rapidly evolving virus strains of SARS-CoV-2. There is a lack of research on broad-spectrum natural compounds that have proven effective in inhibiting coronavirus growth and replication. Many successful drug products are derived from medicinal herbs and then re-engineered to produce the active compound using synthetic chemistry. A broadspectrum viral inhibitor that targeting s various lineages of SARS-CoV-2 by blocking the initial stage of infection, including virus uptake and proliferation, may reduce the infectious viral load and, thereby, disease severity in patients.

We have studied Bi121, an aqueous polyphenolic-rich fraction from *Pelargonium sidoides* (PS). The aqueous ethanolic PS extract has been used as a traditional medicine for the treatment of various ailments for over a century (8). A proprietary extract from PS roots known as EPs7630 or Umckaloabo has been evaluated in numerous clinical trials for safety and alleviation of symptoms associated with acute bronchitis and is licensed in Germany as a medicine for the treatment of upper respiratory tract infections (9). PS extract contains numerous metabolites and was reported to inhibit viruses associated with respiratory diseases like influenza viruses (10–13), HIV (14), and herpes virus (15). Very recently, EPs7630 was shown to have activity against SARS-CoV-2 (16).

More than 30 clinical trials have been conducted with EPs7630 over the last 25 years (total study population > 10,500) in the treatment of acute respiratory tract infections. PS extract has excellent safety and tolerability in human clinical studies (17). This corroborates the safety of Pelargonium-based actives for human applications. Here we investigated the antiviral activity of Bi121 against several strains of SARS-CoV2. The data provide insight into the broad-spectrum antiviral activity of Bi121 against several SARS-CoV2 strains.

### Production and Standardization of Bi121

Crude PS extracts were generated from dried plant roots of PS sourced from South Africa. Roots were ground with a ball mill and 100 g powdered roots were stirred in 600 ml water (ddH_2_O) for 24 h at 55°C. This aqueous extraction was further extracted using ultrasonication (Sonicator 4000, Misonix Inc), and the mixture was centrifuged to get a clear aqueous extract. All PS extract stock solutions were sterilized by filtration and stored at −20°C until used for preliminary functional assays or polyphenolic enrichment.

### Enrichment of polyphenolic-rich Bi121 and fractionation of Bi121 into various fractions

For polyphenolic enrichment, aqueous PS extract was thoroughly mixed with polyvinylpyrrolidone (PVPP, Sigma-Aldrich); polyphenols were allowed to adsorb to PVPP at room temperature for 20 m. The mixture was filtered and washed three times with ddH_2_O. Polyphenols was eluted three times each with 0.5 ml 0.5 N NaOH.

### Bi121 Neutralizes Pseudotyped-SARS-CoV-2 S (spike protein) Representing Various Viral Mutations

We first studied the activity of Bi121 against recombinant Vesicular Stomatitis Virus (rVSV)-pseudotyped SARS-CoV-2 S expressing the firefly luciferase (Integrated BioTherapeutics Inc, Rockville, MD). Briefly, the rVSV with glycoprotein gene (G) deleted was used as the base platform for IBT’s pseudotype-based neutralization assays. The VSV-G glycoprotein was transiently expressed by transfection to produce virus particles in HEK293T cells. To create pseudotyped viruses, the VSV-G was substituted with SARS-CoV-2 Spike protein (Full-length Spike protein lacking terminal eighteen amino acids of the cytoplasmic domain), and the resulting virus - rVSV-ΔG-SARS-CoV-2 S - can be handled at Biosafety level 2 and expresses firefly luciferase. Infection efficiency can be measured by quantifying luciferase activity by reading the relative light units (RLU) on a luminometer. Neutralization activity (IC_50_) of various dilutions of Bi121 against multiple rVSV-ΔG-SARS-CoV-2 S variants expressing spike protein mutations (**Table.1**) were tested in Vero E6 cells. Cytotoxicity of Bi121 (CC_50_) was assessed in parallel.

**Table 1.** Specific mutations of VSV-pseudotyped spike protein that represents various lineages of SARS-CoV-2

| **rVSV-ΔG-SARS-CoV-2 S** | **Specific mutations in the spike protein of pseudotyped strains** |
| --- | --- |
| Wuhan Wild Type | - |
| B.1.1.7 (Alpha) | A570D, D614G, D1118H, delH69-V70, N501Y, P681H, S982A, T716I, delY144 |
| B.1.351(Beta) | K417N, E484K, N501Y, D614G, A701V |
| B.1.617 (Delta) | L452R, E484Q |
| D614G | D614G |
| B.1.427 | L452R, D614G |

For cytotoxicity and neutralization assays, Vero E6 cells were seeded in multiple black 96-well plates 24-hours prior – (Day-1) at 5.00E+04 cells per well with EMEM medium [10% heat-inhibited fetal calf serum (FCS), 2 mM L-Glutamine, 1x Penicillin/Streptomycin (P/S)]. On Day 0, 2-fold dilutions of Bi121 were prepared: For cytotoxicity, all the media were removed from the 96-well plate and replaced with 50 μL of serum-free medium (Gibco, VP-SFM (1x)) supplemented with 2 mM L-Glutamine. 50 μL of Bi121 dilutions were added in triplicate to Vero E6 cells and incubated at 37°C for 24h; medium and cells alone wells were also included as controls. Luciferase activity was measured (Promega, CellTiter-Glo^®^ 2.0 Cell Viability Assay), and Bi121 CC_50_ was determined.

For neutralization, 50 μL of each Bi121 dilutions were mixed with 50 μL rVSV-ΔG-SARS-CoV-2 S variants in a 1:1 ratio for 1hour at 37°C. All the media were removed from the 96-well plates, and the 100 μL mixtures were added in triplicate to Vero E6 cells for 24h incubation at 37°C; pseudotyped virus only and cells only wells were also included as controls. Luciferase activity was measured (Promega, Bright-Glo™ Luciferase Assay System) to determine Bi121 IC_50_. Neutralization assays were validated using Covid-19+ rat serum (data not shown). Data analyses were conducted using XLFit (4 Parameter Logistic Model or Sigmoidal Dose-Response Model). Bi121 effectively neutralized multiple rVSV-ΔG-SARS-CoV-2 S variants expressing different mutations with Bi121 IC_50_ dilutions ranging from 183-1350 (**Figure. 1A-E**). Bi121 IC_50_ values against rVSV-ΔG-SARS-CoV-2 S WT and rVSV-ΔG-SARS-CoV-2 S Delta was 183 (**Figure. 1A**) and 999.6 (**Figure. 1D**), respectively. This preliminary screening showed the broad-spectrum neutralizing ability of Bi121 to rVSV-ΔG-SARS-CoV-2 with various spike protein mutations.

**Figure 1.**
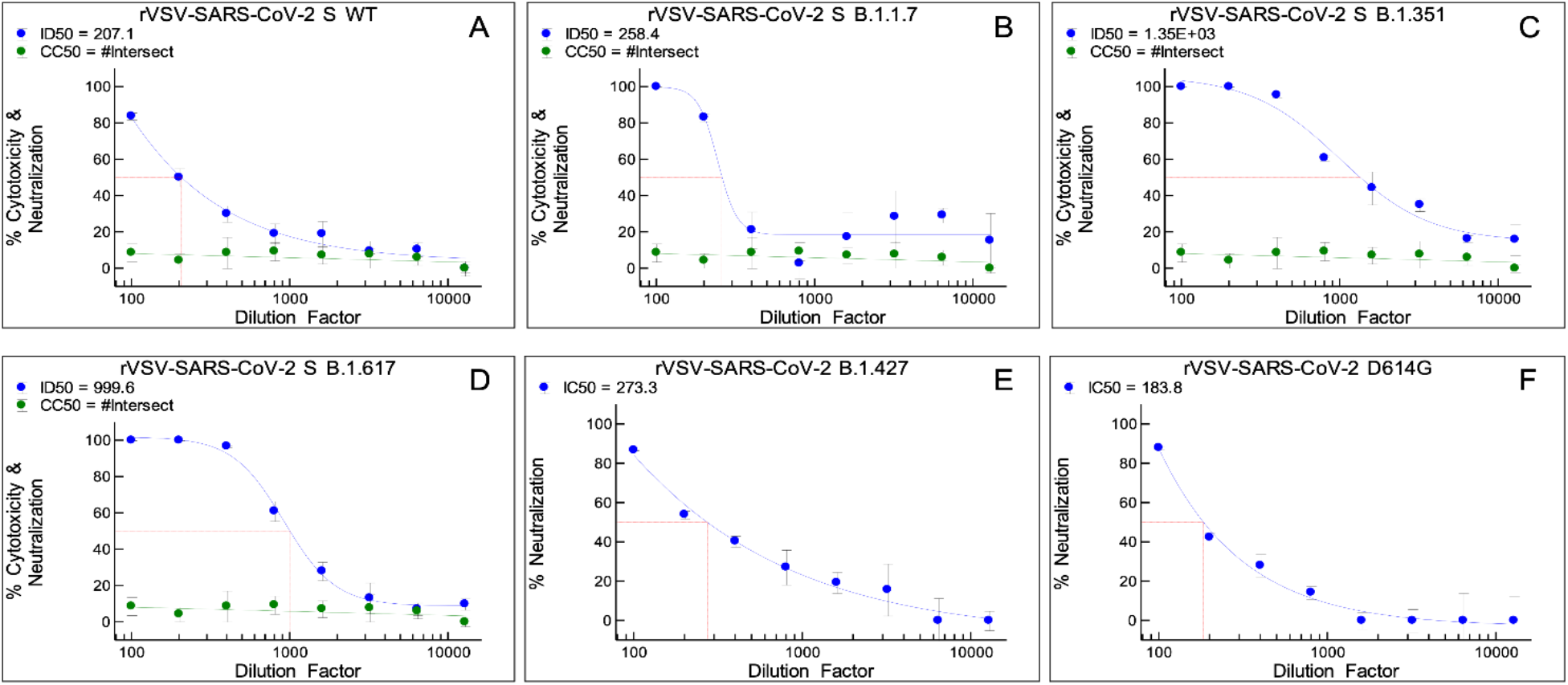
Cytotoxicity (**A-D**) and neutralization activity of Bi121 (**A-F**) against six rVSV-ΔG-SARS-CoV-2 S variants. **A**. rVSV-ΔG-SARS-CoV-2 S WT; **B**. rVSV-ΔG-SARS-CoV-2 S B.1.1.7 **C**. rVSV-ΔG-SARS-CoV-2 S B.1.351 **D**. rVSV-ΔG-SARS-CoV-2 S B.1.617 **E**. rVSV-ΔG-SARS-CoV-2 S B.1.427 **F**. rVSV-ΔG-SARS-CoV-2 S D614G. Cytotoxicity is shown in green and was not assessed for rVSV-ΔG-SARS-CoV-2 S B.1.427 and rVSV-ΔG-SARS-CoV-22 S D614G. Neutralization is shown in blue.

### Bi121 Neutralizes Three Variants of SARS-CoV-2 in Two Different Cell Lines

To test the antiviral activity of Bi121 against SARS-CoV-2, Biom Pharmaceuticals partnered with the Center for Infectious Disease Research at the Indian Institute of Sciences (IISc., Bangalore, India). All experiments entailing live SARS-CoV-2 followed the approved standard operating procedures for viral biosafety level 3 facility approved by the IISc biosafety committee. After observing the cytopathic blocking effect of Bi121 in various dilutions (results not shown), the antiviral activity of Bi121 was assessed against three variants of SARS-CoV-2 using RT-qPCR, plaque assay, and TCID_50_ estimation in two different cell lines (Vero E6 cells and HEK-ACE2). The three SARS-COV-2 variants tested were: USA WA1/2020, Hongkong/VM20001061/2020, B.1.167.2 (Delta).

As shown in **Figure. 2**, Bi121 showed significant activity towards all the three SARS-CoV-2 variants tested in two cell lines. In Vero E6 and HEK-ACE2 cells, Bi121 at 1:40 dilution significantly reduced viral replication (4-5 log reduction) compared to untreated by RT-qPCR, plaque assays, and TCID_50_ estimation for USA-WA1/2020 and Hongkong/VM20001061/2020 strains (**Figure. 2**). Similarly, Bi121 inhibited B.1.167.2 (Delta) replication in Vero E6 and HEK-ACE2 cells by 4-5 log using RT-qPCR and TCID_50_ assay and by 2-3 log using plaque assay (**Figure. 2**).

**Figure 2.**
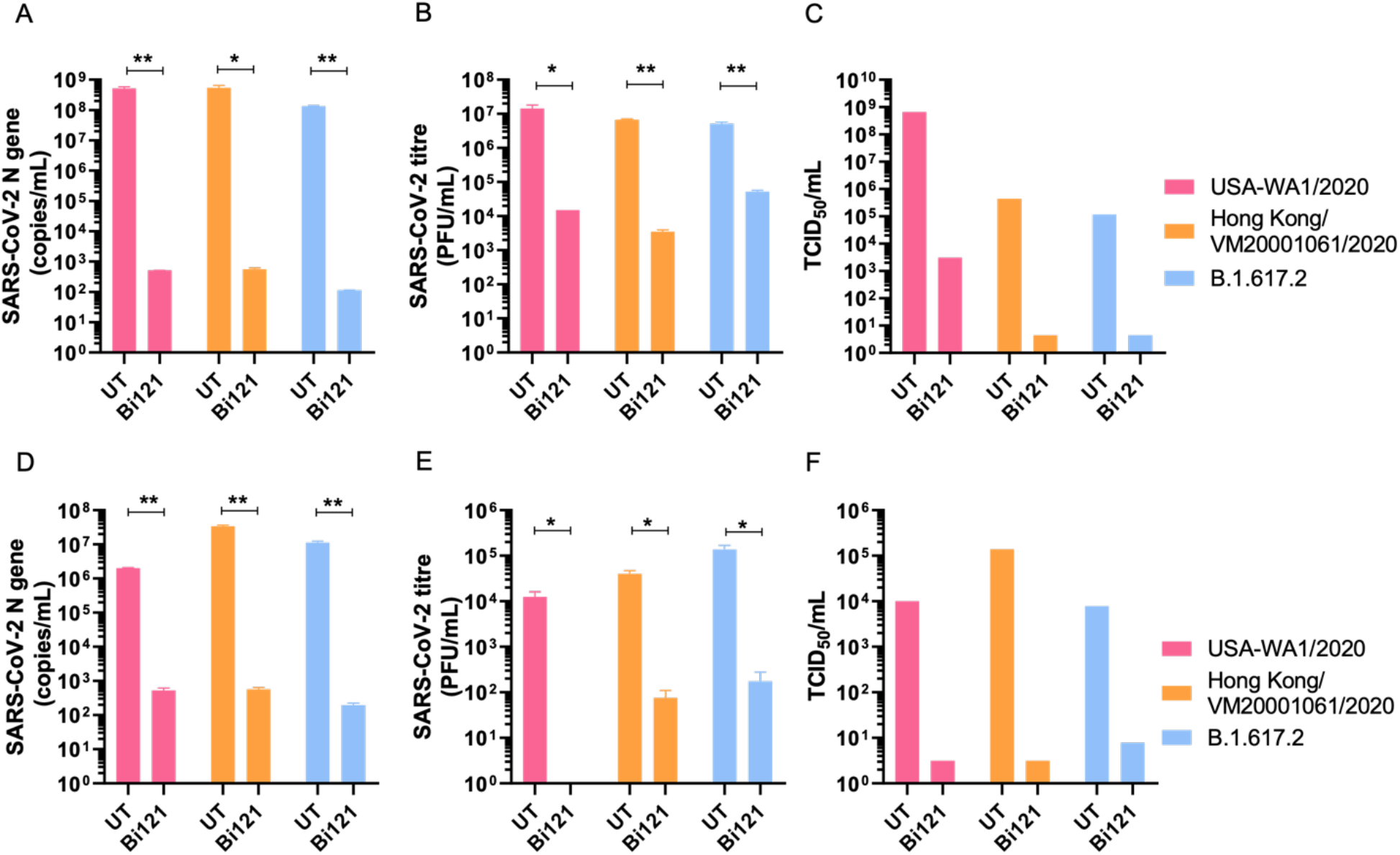
Anti-SARS-CoV-2 activity of Bi121: The antiviral activity of Bi121 was assessed against multiple strains of SARS-CoV-2 by RT-qPCR (**A & D**), plaque assay (**B & E**) and TCID_50_ estimation (**C & F**). Vero E6 (**A, B & C**) and HEK-ACE2 cells (**D, E &, F**) were pre-treated for 2 h with Bi121 (1:40 dilution) or left untreated and were subsequently infected with indicated SARS-CoV-2 strains at a MOI of 0.05 for Vero E6 and 0.1 for HEK-ACE2 cells for 1 h. Viral inoculum were washed, and medium was replaced with fresh media (DMEM with 2% FBS) containing Bi121 or left untreated (UT). Post-infection at 48 h, medium supernatant was harvested and processed for RNA isolation or serially diluted to perform plaque assay and TCID_50_ estimation. SARS-CoV-2 RNA was isolated from 140 μL of supernatant and levels of SARS-CoV-2 specific N-gene was assessed by RT-qPCR. Potential antiviral activity was assessed by comparing compound treated well to untreated and represented as RNA copy numbers. **(A & D).** Infectious SARS-CoV-2 titre were assessed in Vero E6 cells by standard plaque assay and TCID_50_ estimation **(B, C, E & F)**. Error bar represents standard deviation from the mean of two independent experiments (N=2). **, p<0.001; *, p<0.05, by unpaired student’s *t*-test.

### Bi121 Interfere with Early Stages of SARS-CoV-2 Infection

The primary mode of Bi121 action on SARS-CoV-2 replication was evaluated by time-of-drug-addition experiments. HEK-ACE2 cells were either pre-treated or left untreated with Bi121 (1:40 dilution) for 2 h and subsequently infected with SARS-CoV-2 for 1 h. After 48 h post infection, medium supernatants were harvested and processed for RNA isolation and RT-qPCR. The schematic of the experiment is shown in **Figure 3A. Figure. 3B** represents the results obtained as mean values of two independent experiments. Bi121, when added 2 h before and after the infection, showed an inhibitory effect (p=0.0011). Bi121, when added 8 h after infection, showed a less pronounced inhibitory effect (p=0.0027). This suggests that Bi121 may interfere in the early steps of SARS-CoV-2 entry and replication. The inhibitory activity of Bi121 may result from multivalent interactions between the polyphenolic molecules to bind tightly to the S-protein of SARS-CoV-2 to prevent the initiation of viral cell entry and multiplication. This effect was shown with various SARS-CoV-2 strains and multiple mutants, suggesting a broader inhibitory interaction than specific antibody-mediated neutralization.

**Figure 3.**
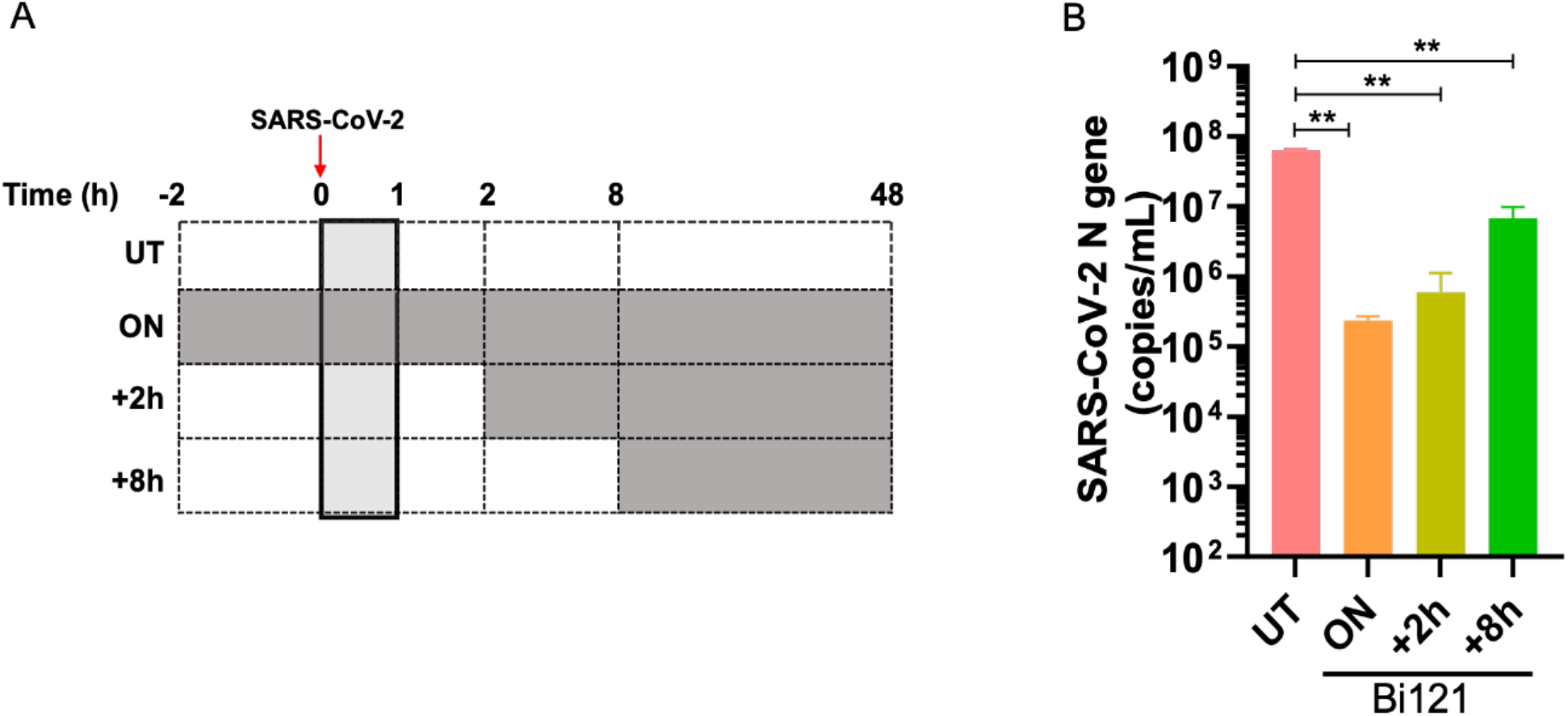
Early treatment with Bi121 inhibits SARS-CoV-2 replication. Time of addition assay was performed to identify the effect of Bi121 on SARS-CoV-2 replication. HEK-ACE2 cells were either pre-treated or left untreated with Bi121 (1:40 dilution) for 2 h and subsequently infected with 0.1 MOI of SARS-CoV-2 (Hongkong/VM20001061/2020) for 1 h. Viral inoculum were washed, and medium was replaced with fresh media (DMEM with 2% FBS) containing Bi121 at indicated time points. Post-infection at 48 h, medium supernatant was harvested and processed for RNA isolation. (A) Schematic of the experiment. Grey boxes represent Bi121 treatment. (B) SARS-CoV-2 RNA was isolated, and levels of SARS-CoV-2 specific N-gene was assessed by RT-qPCR. Potential antiviral activity was assessed by comparing compound treated wells to untreated and represented as RNA copy numbers. Error bar represents standard deviation from the mean of two independent experiments (N=2). **, p<0.001, by unpaired student’s *t*-test.

### Antiviral Activity of Various Fractionated Polyphenolic Compounds from Bi121

To further understand the antiviral activities of Bi121, we fractionated the polyphenol-rich Bi121 into nine fractions (F0-F8) using HPLC for molecular characterization and to identify potential anti SARS-CoV2 compounds. The fractionation of polyphenol enriched fraction was carried out using an Agilent AdvanceBio Column (2.7 μm, 2.1 x 250 mm) at Solvent A (10 mM TEABC, pH 8.0) and an Agilent UHPLC 1290 system. The separation was performed by running a gradient of Solvent B (10 mM TEABC, pH 8.0, 90% ACN) and Solvent A (10 mM TEABC, pH 8.0) at the flow rate 250 μl/min. The eluted fractions were collected according to retention time (12 minutes per fraction) and a total of nine fractions were collected, further evaporated by using speed vacuum. Our preliminary characterization of Bi121 using mass spectroscopy showed that it contains, sesquiterpenes, prodelphinidins and undefined oligomeric carbohydrates (**Figure. 4**).

**Figure 4.**
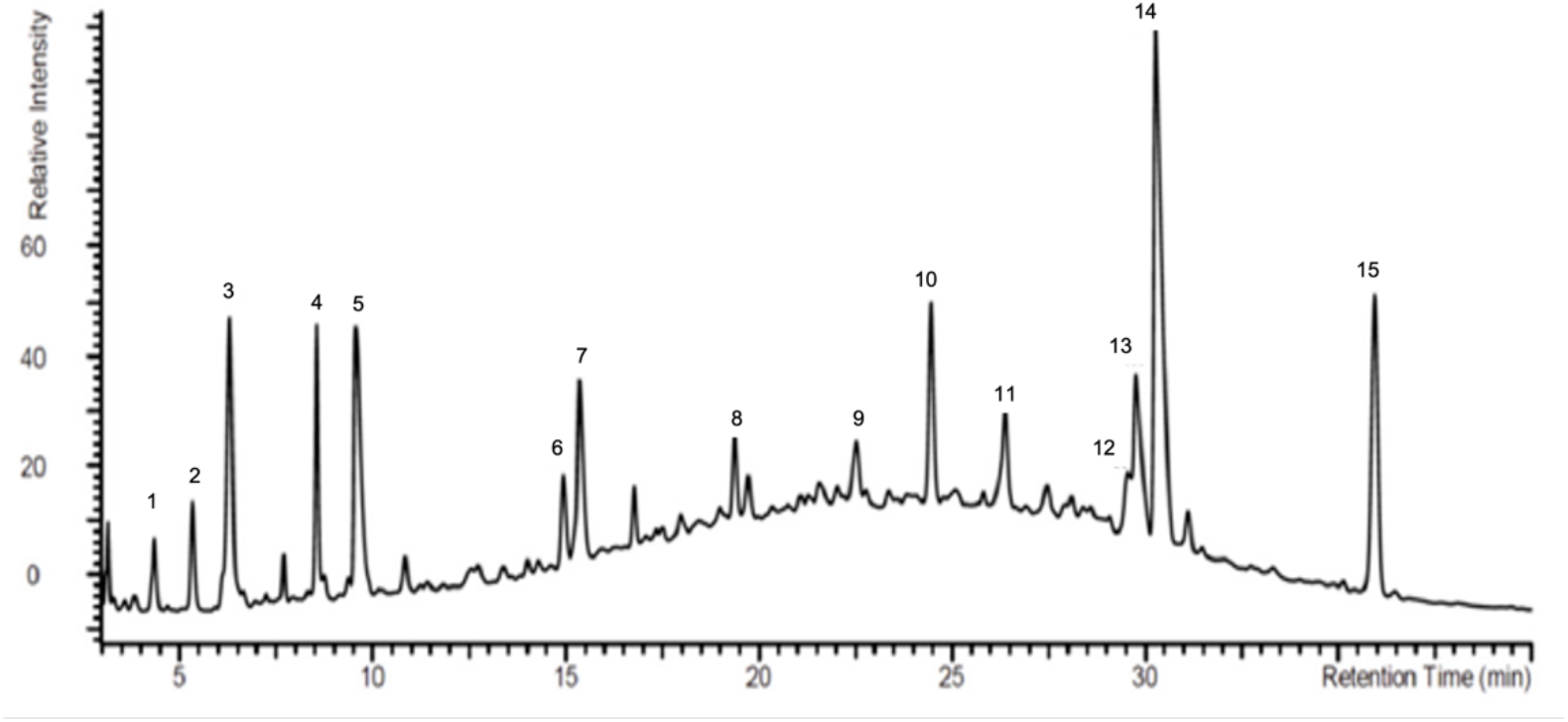
HPLC profile of standardized polyphenolic-rich Bi121 with fifteen characteristic compound peaks.

The antiviral activity was assayed after Vero cells were pretreated with increasing concentration of various fractions for 2 h and subsequently infected with SARS-CoV-2 at an MOI of 0.01. After 48 h post infection, MTT assay was performed to determine the cell viability. Cell viability was calculated after normalizing the data with uninfected control. Percentage viral inhibition was calculated by normalizing the viability of treated cells with respect to viability of untreated controls.

The viral inhibition of various collected fractions was studied using MTT assay. Briefly, Vero cells were pretreated with increasing concentration of each fraction (F0-F-8) for 2 h and subsequently infected with SARS-CoV-2 at an MOI of 0.01. 48h post infection MTT assay was performed to determine the cell viability. Cell viability was calculated after normalizing the data with uninfected control. Percentage viral inhibition was calculated by normalizing the viability of treated cells with respect to viability of untreated controls as follow; % Viral Inhibition = [(cell viability after treatment-cell viability of untreated)/cell viability of untreated]*100. A minimum of 10% viral inhibition was set as cut off value to determine the potential antiviral fraction.

Based on the viral inhibition assays, B121.1, and fractions F2, F3 and F5 showed antiviral activity against SARS-CoV-2 (**Figure. 5**) suggesting multiple compounds with activity against SARS-CoV-2. Fraction 5 showed 40% viral inhibition at 6.25μg/mL, however increasing concentration showed lower viral inhibition. Polyphenolic compounds tend to form polymeric compounds which may be the reasons for lesser activity at higher concentration. Our preliminary assays have shown the predominant compounds responsible for the SARS-CoV-2 antiviral activity in F2, F3, and F5 fractions are sesquiterpenes, and a detailed molecular identification of the compounds in B121 fractions (F2, F3, F5) using mass spectroscopy is underway.

**Figure 5.**
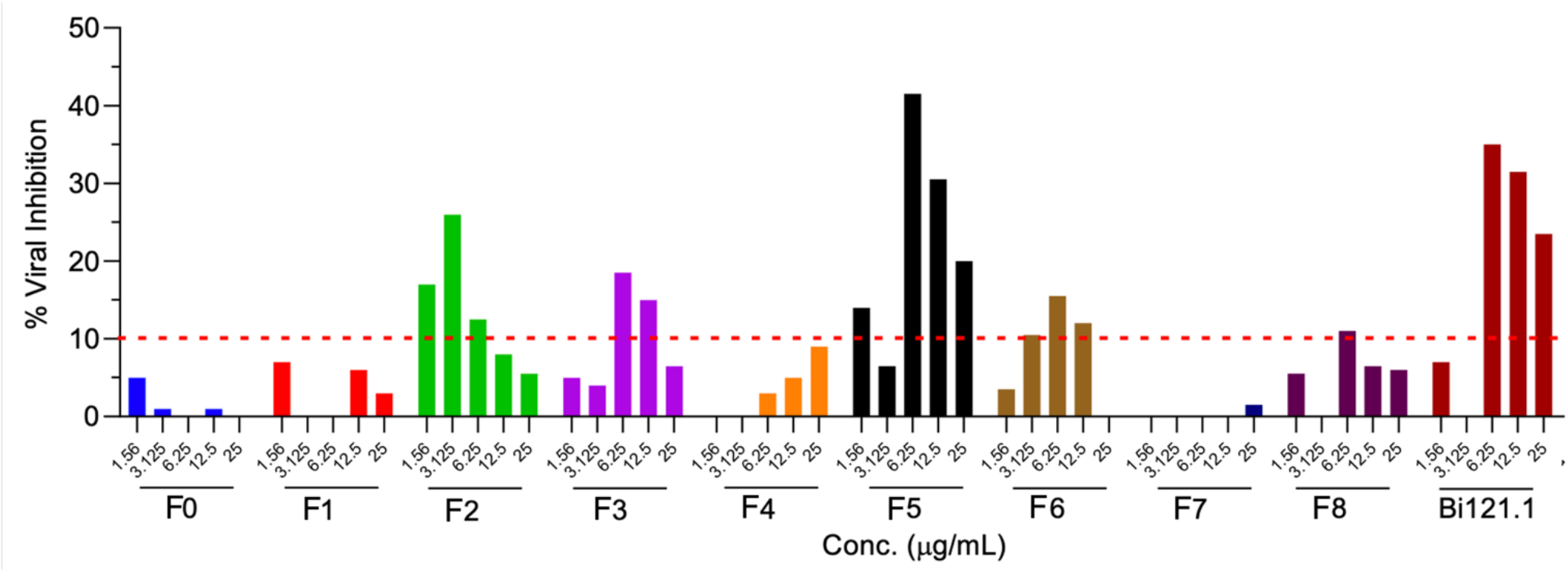
Anti SARS-CoV2 effect of different fractionated compounds. Vero cells were pretreated with indicated concentration of compounds for 2 h and subsequently infected with SARS-CoV-2 at an MOI of 0.01. 48 h post infection MTT assay was performed to determine the cell viability. Cell viability was calculated after normalizing the data with uninfected control. % Viral inhibition was calculated by normalizing the viability of treated cells with respect to viability of untreated controls as follow; % Viral Inhibition= [(cell viability after treatment-cell viability of untreated)/cell viability of untreated] *100. A minimum of 10% viral inhibition was set as cut off value to determine the potential antiviral fraction.

## Summary

In summary, we demonstrated that Bi121 significantly inhibited various strains of SARS-CoV-2 in the two cell lines tested. This inhibition was also observed with the rVSV-ΔG-SARS-CoV-2 system that represents spike protein mutations of various SARS-CoV-2 lineages. Moreover, Bi121 used at concentrations that inhibited the virus did not display toxicity to the target cells. Bi121 treatment before infection or early in the infection showed an inhibitory effect suggesting a treatment as well as a prophylactic measure against SARS-CoV-2. Projected epidemiological consequences of the Omicron SARS-CoV-2 show a rapid increase in infections in the EU and US in the coming months (18). A record number of one million cases per day of new Omicron infections has already been reported in the US on the first week of January 2022. Although preliminary data indicates that Omicron may have less severe infections, the variant is better and faster in transmission (19).

Broad-spectrum antiviral against SARS-CoV-2 is a global need as new virus lineages with more immune-thwarting mutations are evolving (20). Bi121 has the potential to become a successful therapeutic because of the strong antiviral activity without cytotoxic effects and is a thermally stable compound without the need for special logistics support for distribution. This is particularly important for low-resource setting countries that doesn’t have special transportation needs or logistics support. There is significant safety information (8–12) on use of Pelargonium-based actives for human respiratory applications that warrants a fast-track clinical development.

The nasal epithelium is where most respiratory viral infections, including SARS-CoV-2 infects (7). Development of Bi121 in a nasal liquid dosage as a preventative treatment on nasal tissue may reduce virus. We are developing Bi121 as a therapeutic nasal spray for the first line of defense against SARS-CoV2 infection. We are currently testing its efficacy in the MA10 animal model, a SARS-CoV-2 *in vivo* model (21) and identifying antiviral molecular entities in B121 fractions using mass spectroscopy.

## Declaration of Interest

All authors have completed the ICMJE uniform disclosure form at www.icmje.org/coi_disclosure.pdf and declare: AS has received research grants from Biom Pharmaceutical Corporation and Council Of Scientific and Industrial Research, Govt of India to conduct the SARS-CoV-2 research in this article. BS is an executive leadership member at Biom Pharmaceutical Corporations management team. Bi121 formulation and its use against coronaviruses are protected by a US patent application.

